# Benchmarking Sequence-Based and AlphaFold-Based Methods for pMHC-II Binding Core Prediction: Distinct Strengths and Consensus Approaches

**DOI:** 10.1101/2024.10.06.616783

**Authors:** Soobon Ko, Honglan Li, Hongeun Kim, Woong-Hee Shin, Junsu Ko, Yoonjoo Choi

## Abstract

**Background:** Interactions between peptide and MHC class II (pMHC-II) are crucial for T-cell recognition and immune responses, as MHC-II molecules present peptide fragments to T cells, enabling the distinction between self and non-self antigens. Accurately predicting the pMHC-II binding core is particularly important because it provides insights into pMHC-II interactions and T-cell receptor engagement. Given the high polymorphism and peptide-binding promiscuity of MHC-II molecules, computational prediction methods are essential for understanding pMHC-II interactions. While sequence-based methods are widely used, recent advances in AlphaFold-based structure prediction have opened new possibilities for improving pMHC-II binding core predictions.

**Results:** We benchmarked four recent pMHC-II prediction methods with a focus on binding core prediction: two sequence-based methods, NetMHCIIpan and DeepMHCII, and two AlphaFold-based structure prediction methods, AlphaFold2 fine-tuned for peptide interactions (AF2-FT) and AlphaFold3 (AF3). The AlphaFold-based methods showed strong performance in predicting positive binders, with AF3 achieving the highest positive recall (0.86) and AF2-FT performing similarly (0.81). However, both methods frequently misclassified unbound peptides as binders. NetMHCIIpan excelled at identifying non-binders, achieving the highest negative recall (0.93), but had lower positive recall (0.44). In contrast, DeepMHCII demonstrated moderate performance without any notable strength. Consensus approaches combining AlphaFold-based methods for binder identification with filtering using NetMHCIIpan improved overall prediction precision (0.94 and 0.87 for known and unknown binding status, respectively).

**Conclusions:** This study highlights the complementary strengths of AlphaFold-based and sequence-based methods for predicting pMHC-II binding core regions. AlphaFold-based methods excel in predicting positive binders, while NetMHCIIpan is highly effective at identifying non-binders. Future research should focus on improving the prediction of unbound peptides for AlphaFold-based models. Since NetMHCIIpan’s binding core predictive ability is already high, future efforts should concentrate on enhancing its binding prediction to further improve overall accuracy.

## Background

Antigen presentation is a critical process in the immune response, where antigen-presenting cells, such as dendritic cells, macrophages, and B cells, display peptide fragments on their surface bound to major histocompatibility complex (MHC) molecules. This interaction is pivotal for the activation of T cells, which recognize the presented peptides via their T-cell receptors (TCRs). The process is essential for distinguishing between self and non-self antigens, enabling the immune system to mount responses against a wide variety of pathogenic invaders [1]. Therefore, an accurate understanding of peptide-MHC (pMHC) interactions is crucial, not only for elucidating the mechanisms of immune responses but also for the development of effective vaccines and immunotherapies [2–6].

MHC molecules are highly polymorphic [7, 8], possibly as an evolutionary strategy to protect against the virtually limitless number of potential pathogens [9, 10]. This extreme polymorphism allows MHC molecules to present a broad spectrum of antigenic peptides. In addition to the polymorphism, a single MHC molecule can bind to a variety of different peptides [11]. This broad peptide-binding capacity, or peptide-binding promiscuity, further enhances the immune system’s flexibility, allowing it to respond to many different pathogens despite the limited number of MHC alleles in any given individual. However, the polymorphism and peptide-binding promiscuity of MHC molecules make it practically impossible to explore the entire pMHC interaction space experimentally, presenting a significant challenge for immunological research [12, 13].

Given the impracticality of experimentally determining all pMHC interactions, computational prediction methods have emerged as essential tools for investigating these interactions. There are two main approaches for predicting peptide-MHC interactions: sequence-based and structure-based methods [14, 15]. Traditionally, sequence-based methods have been more widely used due to their speed and accuracy. These methods rely on large datasets of known pMHC binding affinities to train machine learning models [16–19] or apply position-specific scoring matrices [20–22]. Sequence-based methods are generally more effective for predicting pMHC-I interactions, where the peptide length is relatively fixed (typically 8-11 amino acids). This relatively fixed length allows for more straightforward modeling of the binding core region, leading to high prediction accuracy. However, predicting pMHC-II interactions is more challenging. Unlike MHC-I, MHC-II molecules bind longer peptides with additional flanking regions at each terminus, making it difficult to accurately identify the binding core region. As a result, sequence-based methods often struggle with the greater flexibility of pMHC-II complexes [23–25].

Structure-based approaches have gained increasing attention due to recent breakthroughs in artificial intelligence and its application to protein structure prediction. Tools such as AlphaFold [26] and RoseTTAFold [27] have revolutionized the field, enabling the reliable prediction of protein structures with remarkable accuracy. However, structure-based methods have been less effective in modeling immune molecules, primarily because co-evolutionary information is often unavailable for these interactions [28]. To address this limitation, the development of AlphaFold2 fine-tuned for peptide interactions has shown promising results [29]. Additionally, the most recent version, AlphaFold3 [30], has demonstrated enhanced prediction accuracy, delivering impressive performance in molecular modeling.

In this study, we benchmarked two widely-used sequence-based methods, NetMHCIIpan [31] and DeepMHCII [32], and two structure-based methods, AlphaFold2 fine-tuned for peptide interactions [29] and AlphaFold3 [30], with a specific focus on identifying the binding core regions (**Fig. 1A**). While there have been numerous benchmark and performance comparison studies for pMHC-II prediction, to date, no studies have specifically focused on the accuracy of binding core region prediction.

**Figure 1.**
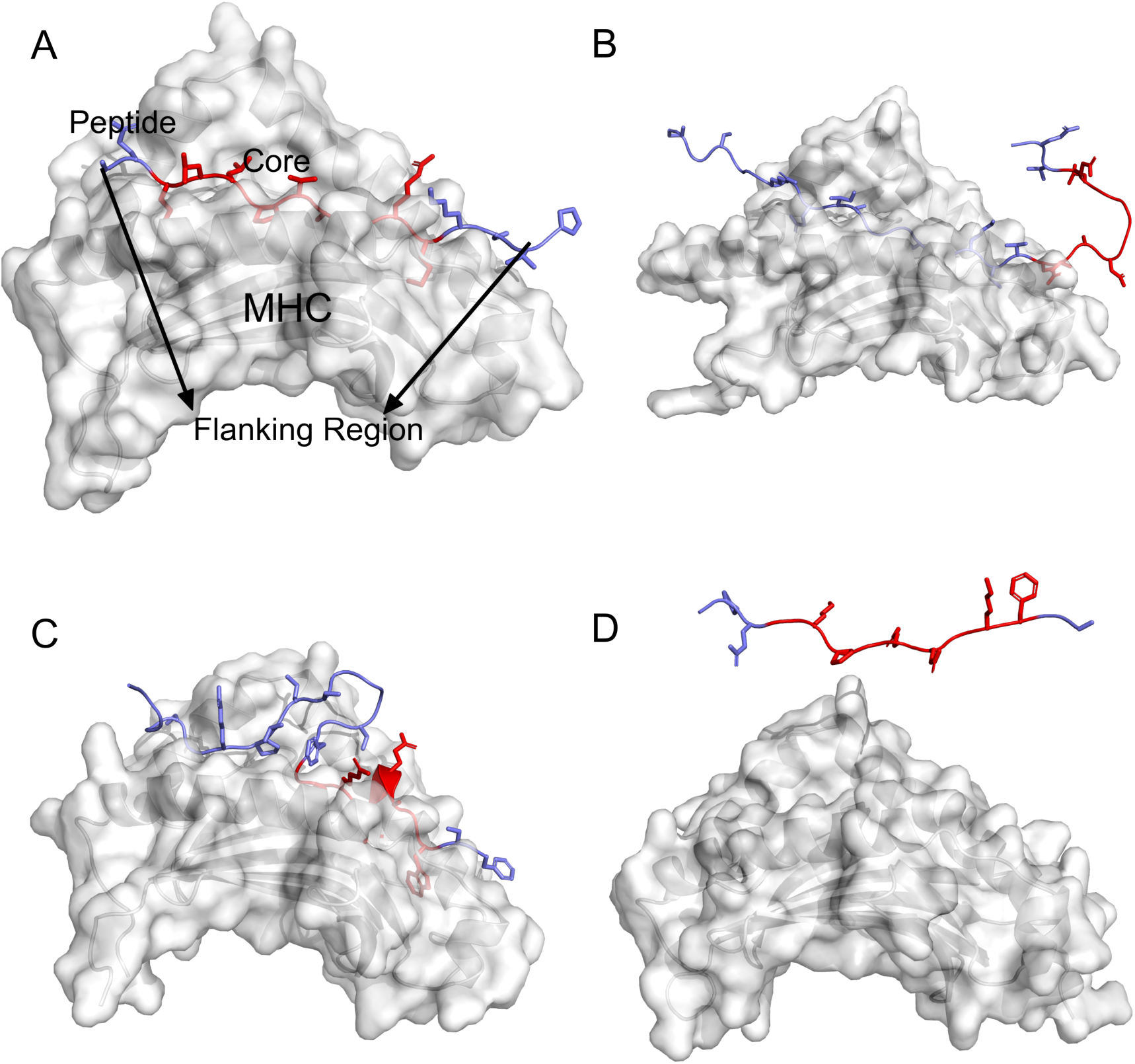
Examples of pMHC-II binding and prediction assessment. The MHC-II complex is shown in white, the non-core portion of the peptide is colored in blue, and the core region is in red.(A) The pMHC-II complex with the correctly predicted core region. (B) The peptide binds to MHC- II, but the predicted core region is incorrect. (C) Incorrect peptide binding conformation is regarded as unbound. (D) Complete mislocation of the peptide is also considered unbound.

To address this gap, we prepared an extensive test set and evaluated the computational prediction methods in various settings, providing a comprehensive comparison. Our results show that each method has distinct characteristics in prediction. We propose a consensus approach that combines the strengths of both sequence-based and structure-based methods to improve overall prediction accuracy and provide more reliable results for pMHC-II binding core region identification.

## Methods

### Dataset preparation

A total of 126 peptide-MHC-II (pMHC-II) complex structures, including those with T-cell receptors, were collected from the IMGT database [33]. Any structures containing non-standard amino acids within the pMHC-II were excluded. For the MHC-II structure, only the groove region, as defined by the IMGT, was used for structural modeling [34]. The MHC types and sequences were identified based on IMGT annotations [35]. From this initial set, a non-redundant set of 72 unique sequences was prepared for benchmarking (**Supplementary File 1**). Unbound peptides were generated by randomly shuffling the positive peptides (the same length).

### Computational prediction methods and assessment of binding

To benchmark predictions of the binding core region in pMHC-II complexes, we employed two sequence-based methods (NetMHCIIpan-4.3 [31] and DeepMHCII [32]) alongside two structure-based methods, AlphaFold2 finetuned for peptide binding (AF2-FT)[29] and AlphaFold3 (AF3)[30]. The default settings for each prediction tool were used, and binding predictions were assessed based on specific criteria for each method.

For NetMHCIIpan-4.3, a 5% rank threshold was applied to identify binders, which included both weak and strong binders. DeepMHCII classified peptides as binders if their predicted IC50 binding values were below 500 nM. For the two AlphaFold-based prediction methods, top-ranked predicted structures were used to estimate pMHC-II binding. All AF3 predictions were performed using the webserver (https://alphafoldserver.com/). For AF2-FT, the query peptide sequence consisted of 11 residues, with a core nine-residue segment and one residue at each terminus. For peptides longer than 11 residues, overlapping windows were generated by sliding the window along the peptide sequence. The top four templates were selected based on MHC alignment scores from a template search and used to construct structural features for AlphaFold modeling. For the selection of final structures, the predicted aligned error (PAE) scores were used, with the window slide showing the lowest PAE being chosen for further analysis of binding and core prediction. In the structure-based methods, binding was assessed manually by inspecting the orientation of peptide side chains relative to the target crystal structure. Predictions were classified as non-binding if side chain orientations in the peptide core region were not aligned with the reference structure (e.g., **Fig. 1C** and **1D**).

## Results

### Prediction performance of pMHC-II binding

We first benchmarked the predictive abilities of the four computational prediction methods, AlphaFold3 (AF3) and AlphaFold2 fine-tuned for peptide interactions (AF2-FT), alongside two sequence-based methods, NetMHCIIpan and DeepMHCII, for peptide binding (without considering true binding cores). The computational methods were tested on a non-redundant set of 72 pMHC-II complexes with determined crystal structures.

The structure-based methods excelled in predicting positive binders (**Fig. 2** and **Table 1**). AF3 achieved the highest positive recall (0.88), reflecting its ability to correctly identify the majority of true binders. AF2-FT performed similarly well, with both methods producing comparable positive F1 scores (AF3: 0.75, AF2-FT: 0.73), demonstrating strong performance in capturing pMHC-II binding events. However, both models exhibited a tendency to misclassify unbound peptides as binders, as indicated by their relatively low negative recall values (AF3: 0.56, AF2-FT: 0.63). This behavior likely stems from the nature of AlphaFold-based structure-based modeling, where the method tends to put peptides into the MHC groove, even when a perfect binding fit is not available [36].

**Figure 2.**
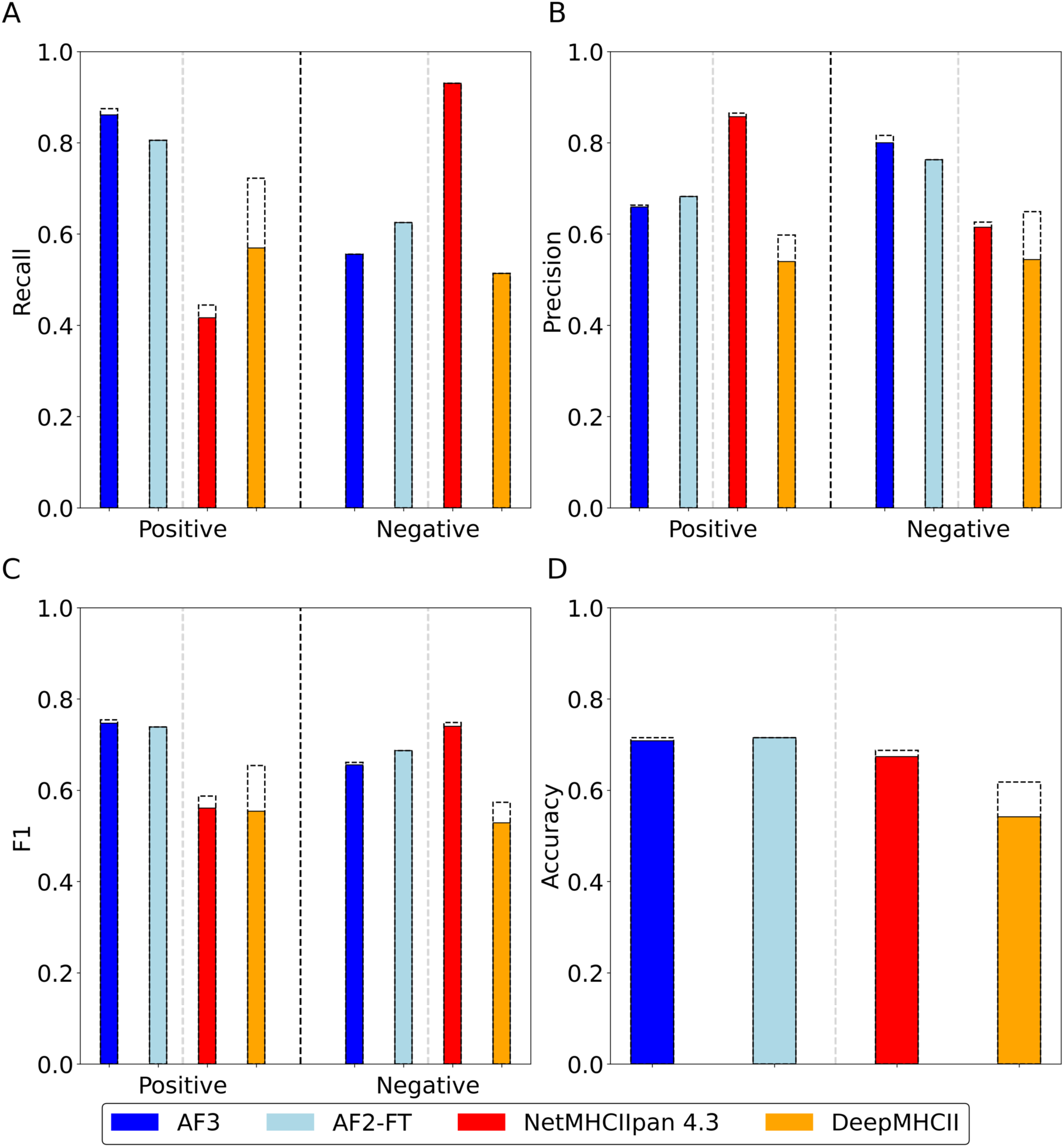
Performance comparison of pMHC-II core prediction methods. The colors represent the different prediction methods: blue for AF3, sky blue for AF2-FT, red for NetMHCIIpan, and orange for DeepMHCII. The dotted bars indicate the performance for binding prediction. When AF2-FT correctly predicts peptide binding, the binding core is always accurately identified, while AF3 misses only one case. In contrast, the sequence-based methods often incorrectly predict core regions even when pMHC-II binding is correctly predicted.

**Table 1.** pMHC-II binding and core region prediction results. Values in brackets represent binding prediction results only. The discrepancies between peptide binding prediction and core prediction are more pronounced in sequence-based methods compared to structure-based methods. Notably, AF2-FT shows no differences between binding and core prediction.

|  | AF3 | AF2-FT | NetMHCIIpan | DeepMHCII |
| --- | --- | --- | --- | --- |
| **Positive Prediction (Binder Prediction)** |  |  |  |  |
| **Recall** | <u>**0.861**</u><br>**(0.875)** | 0.806 | 0.417 (0.444) | 0.569 (0.722) |
| **Precision** | 0.659<br>**(0.663)** | 0.682 | <u>**0.857**</u> **(0.865)** | 0.539 (0.598) |
| **F1** | <u>**0.747**</u><br>**(0.754)** | 0.739 | 0.561 (0.587) | 0.554 (0.654) |
| **Negative Prediction (Non-binder prediction)** |  |  |  |  |
| **Recall** | 0.556 | 0.625 | <u>**0.931**</u> | 0.514 |
| **Precision** | <u>**0.8**</u> **(0.816)** | 0.763 | 0.615 (0.626) | 0.544 (0.649) |
| **F1** | 0.656<br>**(0.661)** | 0.687 | <u>**0.74**</u> **(0.749)** | 0.529 (0.574) |
| **Overall Prediction** |  |  |  |  |
| **Accuracy** | 0.708<br>**(0.715)** | <u>**0.715**</u> | 0.674 (0.688) | 0.542 (0.618) |
| **MCC** | 0.438<br>**(0.454)** | <u>**0.438**</u> | 0.405 (0.429) | 0.083 (0.241) |

In contrast, NetMHCIIpan demonstrated superior performance in identifying non-binders. It achieved the highest negative recall (0.93) and a strong negative F1 score (0.75), indicating its strength in predicting non-binding peptides. Additionally, NetMHCIIpan’s high positive precision (0.86) suggests that when it predicts a peptide as a binder, the prediction is generally accurate. However, its low positive recall (0.44) indicates that it may overlook some true binders, likely due to its strict criteria for determining binding. DeepMHCII, while performing moderately across most metrics, was the least accurate method overall, with an accuracy of 0.62 and an MCC of 0.24. Its balanced performance, though lower compared to the other methods, suggests that it lacks the distinctive strengths of the other models, particularly in accurately predicting either binders or non-binders.

The results indicate that the structure-based methods are particularly good at identifying positive binders but tend to overestimate binding by misclassifying non-binders. On the other hand, NetMHCIIpan excels at correctly identifying non-binders while maintaining high precision for the binders. Combining these two approaches, leveraging the structure-based methods’ high positive recall values and NetMHCIIpan’s high precision, could provide a synergistic solution, improving overall prediction accuracy and reliability.

### Prediction performance of pMHC-II binding core

We next benchmarked the prediction methods for the identification of the binding core region in pMHC-II complexes. As with the binding prediction assessment, AF2-FT and AF3 demonstrated strong performance in binding core prediction (**Fig. 2** and **Table 1**). AF3 achieved a positive recall of 0.86 (62/72), slightly better than AF2-FT (0.81, 58/72) in overall core identification. Both models showed comparable positive F1 scores (AF3: 0.75 and AF2-FT: 0.74), indicating their similar predictive abilities in capturing binding core events.

Notably, for AF2-FT, whenever it predicts a peptide as a binder, the binding core is always correctly identified. A similar trend is observed for AF3, with only one case where the binding core is incorrectly predicted. In the case of unbound peptides, AF2-FT performed slightly better than AF3 (negative recall 0.69 and 0.66, respectively). The two sequence-based methods showed relatively lower performance in predicting pMHC-II binding core, with recall values roughly half of those seen with the structure-based methods. While DeepMHCII generally underperformed in both binder and non-binder prediction, as reflected in its MCC value (0.08), NetMHCIIpan was notably effective at capturing non-binders, achieving a negative recall of 0.93.

Similar to the binding prediction results, the structure-based methods (AF3 and AF2-FT) and NetMHCIIpan demonstrated distinct strengths in different aspects of binding core prediction. The structure-based methods were highly capable of identifying positive binders, but they frequently misclassified non-binding cores as binding. In contrast, NetMHCIIpan’s ability to accurately identify non-binders makes it a reliable method for negative prediction. Given the complementary strengths of the structure-based methods and NetMHCIIpan, a consensus approach combining the high positive recall of AF3 and AF2-FT with NetMHCIIpan’s superior negative recall and positive precision could provide a more accurate and reliable method for predicting the pMHC-II binding core region.

To investigate the failure cases, we further analyzed the prediction results. The first indicator is the query peptide length. The average peptide length of our test set was approximately 17 residues. For AF2-FT and AF3, the average length of failure cases was around 21 residues (23 and 19 residues for NetMHCIIpan and DeepMHCII, respectively). These results suggest that longer peptides are more challenging to predict accurately. **Figure 3A** illustrates the relationship between query peptide length and misprediction rate.

**Figure 3.**
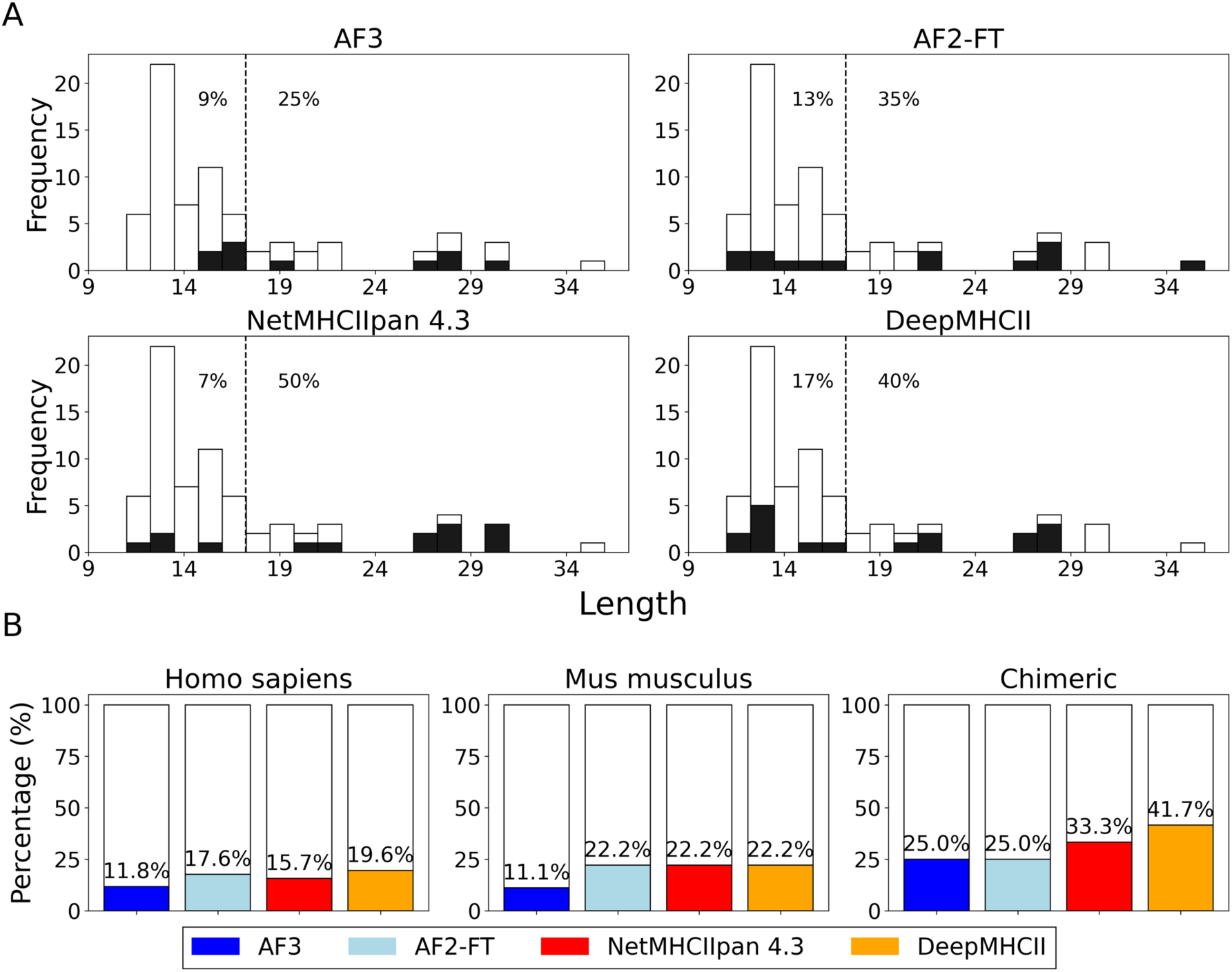
Failure analysis of pMHC-II binding core prediction by peptide length and MHC species. (A) Relationship between query peptide length and misprediction rate for pMHC-II binding core prediction. The average peptide length of the test set was approximately 17 residues, and longer peptides posed a greater challenge for accurate prediction across all methods. (B) Prediction accuracy for different MHC species. Prediction accuracy was significantly lower for peptides bound to chimeric MHCs, particularly for sequence-based methods.

Using 17 residues as a cut-off value for distinguishing long and short peptides, the misprediction rate of AF3 was 9% for short peptides, but increased to 25% for longer peptides. Similarly, AF2-FT recorded 13% for short peptides, which rose to 35% for longer ones. This increase in misprediction rates was even more pronounced for the sequence-based methods. NetMHCIIpan showed a misprediction rate of 7% for short peptides, but the rate increased significantly to 50% for longer ones (For DeepMHCII, 17% and 40% for short and long peptides respectively).

The species of the MHC molecule also plays a significant role in prediction accuracy. Specifically, the prediction accuracy was noticeably lower for peptides bound to chimeric MHCs (**Fig. 3B**). This trend was particularly pronounced for the sequence-based methods, likely due to their inability to effectively handle non-standard MHCs. In contrast, the structure-based methods were relatively less affected by the presence of chimeric MHCs, possibly because they were either trained on a broader dataset or are less data-dependent.

### Consensus approach for pMHC-II binding core prediction

Considering the superior recall of the structure-based methods, we explored two possible scenarios for pMHC-II binding core prediction. In the first scenario, assuming the binding status is experimentally known, we evaluated the performance of each method in predicting the correct core region (**Fig. 4A** and **Table 2**). The structure-based methods showed superior overall performance. Among the 72 binding cases, AF3 correctly predicted the core in 62 cases (86.1%), AF2-FT in 58 (80.6%), NetMHCIIpan in 30 (41.7%), and DeepMHCII in 41 (56.9%). Interestingly, there were several instances where AF3 and AF2-FT did not agree on the predicted core region. By combining the predictions of AF3 and AF2-FT, we obtained the best results, with 68 out of 72 (94.4%) cores correctly predicted.

**Figure 4.**
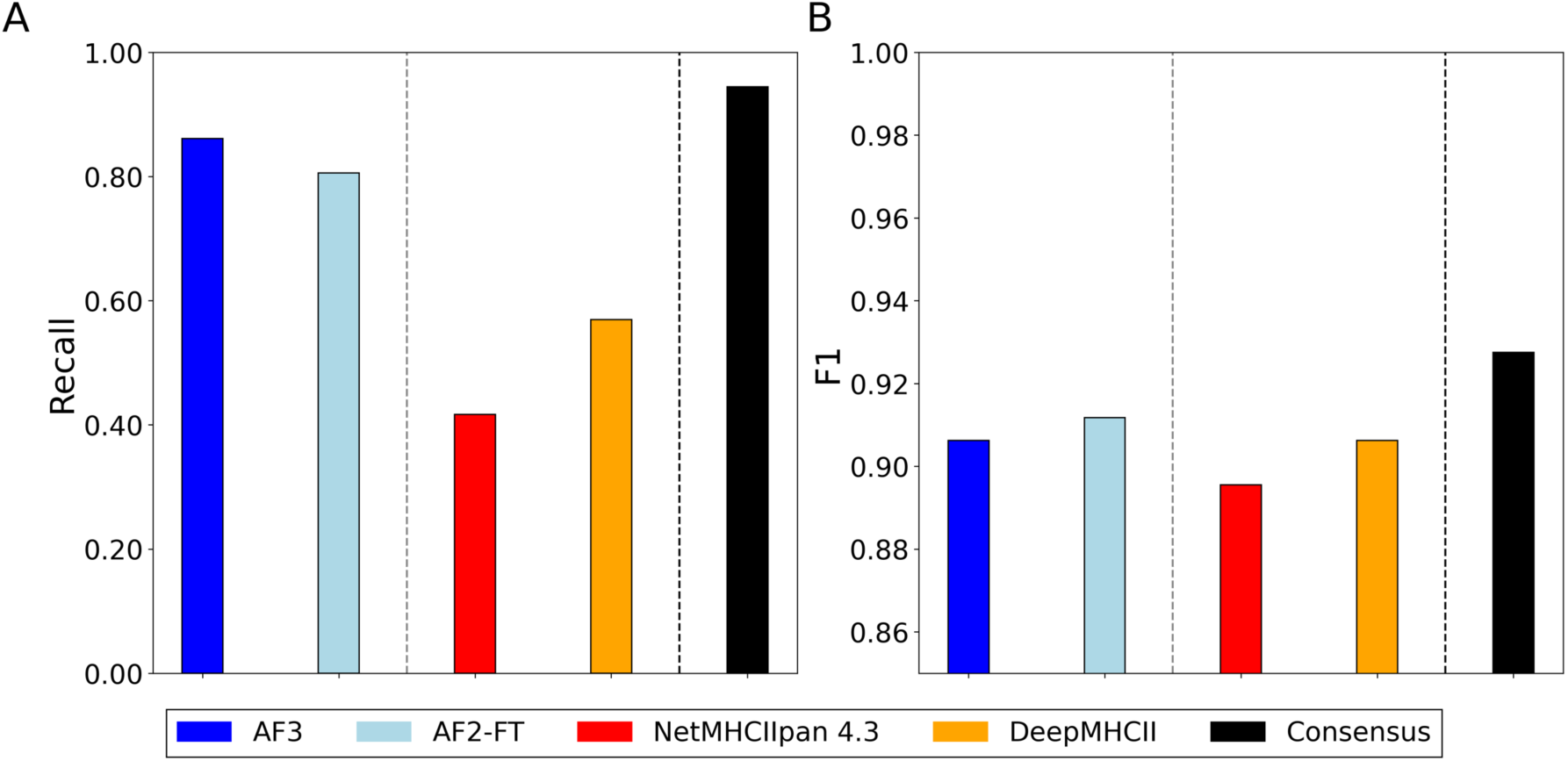
Performance of pMHC-II binding core prediction using consensus approaches. (A) Binding core prediction performance for each method when the binding status is experimentally known. The structure-based methods (AF3 and AF2-FT) demonstrated superior accuracy, with a union consensus approach achieving the highest correct core prediction rate of 94.4%. (B) Binding core prediction performance when the binding status is completely unknown. The combination of AF3 and AF2-FT with NetMHCIIpan filtering further improved precision to 86.5%. The consensus approach demonstrated perfect recall (1.00) and the highest F1 score (0.93).

**Table 2.** Performance metrics of pMHC-II binding core prediction with known vs. unknown binding status.

|  | AF3 | AF2-FT | NetMHCIIpan | DeepMHCII | Consensus |
| --- | --- | --- | --- | --- | --- |
| **Binding status known** |  |  |  |  |  |
| **Precision** | 0.861 | 0.806 | 0.417 | 0.569 | <u>**0.944**</u> |
| **Binding status unknown** |  |  |  |  |  |
| **Recall** | 0.906 | 0.969 | 0.938 | 0.906 | <u>**1.000**</u> |
| **Precision** | <u>**0.906**</u> | 0.861 | 0.857 | <u>**0.906**</u> | 0.865 |
| **F1** | 0.906 | 0.912 | 0.896 | 0.906 | <u>**0.928**</u> |

We further investigated the four failure cases. Notably, two core peptides (WGAEGQRPG for 3T0E and 3O6F, and RLYLVCGEE for 6DFS and 6DFW) were consistently incorrectly predicted. In the former case, the actual query peptide was FS**<u>WGAEGQRPG</u>**FGSGGGSLVPRGSGGGG, and the sequence-based methods agreed on the core region being FGSGGGSLV. This suggests that the sequence-based methods identified a 9mer substring whose termini are hydrophobic, possibly favoring the selection of this region. In the latter case (6DFS and 6DFW), the MHC-II molecule was a chimera between human and mouse, which may explain the failure, as chimeric MHC molecules were likely not included in the training sets of the prediction methods.

We also tested the scenario where the binding status was completely unknown. We first applied a union combination of AF3 and AF2-FT, given their promising performance in the previous setting. This combined approach correctly identified 94.4% of the binders (**Fig. 4A**) but also misclassified 31 out of 72 unbound peptides as binders. Considering that NetMHCIIpan had the highest recall for unbound peptides, we applied a filtering step using NetMHCIIpan to exclude negatively predicted peptides. After filtering, 37 peptides remained, of which 32 were correctly identified, resulting in a precision of approximately 86.5%.

The consensus approach effectively improved overall performance metrics, as shown in **Fig. 4B** and **Table 2**. The consensus method achieved perfect recall (1.00) for the remaining binders, demonstrating its capability to avoid missing any true positive binding cores among the filtered peptides. The F1 score for the consensus approach was also the highest (0.93), highlighting the benefits of integrating complementary strengths to enhance predictive accuracy and reliability.

## Discussion

This study highlights the strengths and limitations of cutting-edge sequence-based and structure-based methods for pMHC-II binding core prediction. While structure-based methods demonstrate high effectiveness in predicting positive binders with correct core identification, they tend to struggle with accurately identifying non-binders. Specifically, these methods often “force” peptides into the MHC groove, leading to an overestimation of binding events. In contrast, sequence-based methods like NetMHCIIpan excel at filtering non-binders but frequently miss true binders. DeepMHCII showed moderate performance across the board, lacking the pronounced strengths seen in the other methods.

The AlphaFold-based structure-based methods tested here were clearly superior in predicting pMHC-II binding cores. AF3, with the highest positive recall and F1 scores, consistently identified most true binders and correct core regions, while AF2-FT performed similarly well. Interestingly, AF3 achieved this high accuracy without any fine-tuning for peptide-MHC interactions, demonstrating the powerful predictive potential of the advanced prediction method. However, their relatively low negative recall suggests that they often misclassify unbound peptides as binders, likely due to the inherent nature of AlphaFold-based modeling that fits peptides into the MHC groove even when the fit is imperfect.

NetMHCIIpan, on the other hand, performed exceptionally well in predicting the binding core region if peptide binding is not considered, with an accuracy comparable to that of AF2-FT (58 out of 72 correct predictions). This suggests that NetMHCIIpan already possesses a strong capability for identifying the correct core region in pMHC-II complexes. However, while its core-binding predictions are generally highly accurate, the method falls short when it comes to predicting whether a peptide will bind at all. This gap shows the need for improvement in its binding prediction capabilities, even though its core identification abilities are already quite advanced.

It is also important to note that all the methods examined in this study are relatively recent, and it remains unclear how much information was already included in their training sets. This uncertainty may impact the evaluation of their performance, especially if some peptides or MHC molecules in our test set were part of their training data, potentially inflating the predictive accuracy. Therefore, future benchmarking studies should consider the training set overlap to ensure a more unbiased assessment.

The consensus approach proposed in this study, which combines the strengths of both structure- and sequence-based methods, provided a synergistic solution to improve overall prediction accuracy. By leveraging the high positive recall of AF3 and AF2-FT for binder prediction and the superior ability of NetMHCIIpan to filter non-binders, we were able to achieve more reliable predictions of pMHC-II binding core regions. This integration of methodologies emphasizes the importance of using a combined approach to balance the strengths of each method.

Looking ahead, future research should focus on addressing the weaknesses of structure-based methods, particularly their tendency to overestimate binding by misclassifying unbound peptides. Enhancing their ability to accurately filter out non-binders will be crucial for improving overall accuracy. Meanwhile, sequence-based methods like NetMHCIIpan would benefit from optimizations that improve binding prediction without sacrificing their core prediction accuracy. Striking this balance between precision and recall will be essential as the field continues to advance.

## Acknowledgements

YC was supported by Medical Technology Development Program of the National Research Foundation of Korea (NRF), funded by the Korean government (MSIT) (2020M3A9G3080281 and 2020R1A5A2031185). WHS acknowledge the support from Ministry of Science and ICT (MSIT), Korea, under the ICAN (ICT Challenge and Advanced Network of HRD) support programs (IITP- 2024-RS-2022-00156439 and IITP-2024-RS-2024-00438263) supervised by the Institute for Information & Communications Technology Planning & Evaluation (IITP). JK and HL were supported by NRF- 2024M3A9J4006525 and SK was supported by RS-2024-00462620.

